# Computational Insights into the Molecular Dynamics of the Binding of Ligands in the Methanol Dehydrogenase

**DOI:** 10.1101/2024.07.08.602606

**Authors:** One-Sun Lee, Sung Haeng Lee

## Abstract

Methanol dehydrogenase (MDH) is a promising biocatalyst for industrial use, converting methanol to formaldehyde. Our molecular modeling revealed methanol binds to MDH with ∼7 kcal/mol free energy, while formaldehyde binds with ∼4 kcal/mol. This suggests methanol remains longer in the active site, and formaldehyde exits more readily post-reaction. These insights are crucial for designing more efficient MDH variants for industrial applications.

**Graphical abstract:** 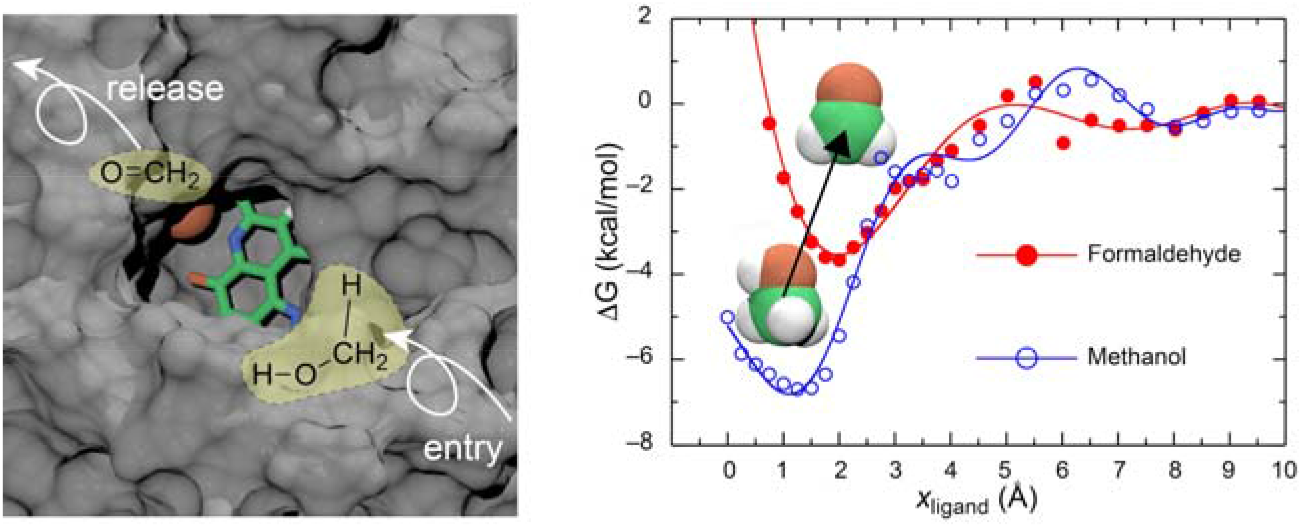

The left panel shows methanol and formaldehyde within the active site of methanol dehydrogenase, illustrating entry and release paths. The right panel presents the free energy profiles, indicating methanol’s stronger binding affinity and formaldehyde’s easier release post-reaction.

---

Methanol dehydrogenase (MDH) is an enzyme in methanotrophic and methylotrophic bacteria that catalyzes methanol oxidation and converts methanol to formaldehyde. In addition to its biological interest, MDH has been regarded as an attractive biocatalyst for industrial applications. Understanding the structure and function of MDH is crucial for its application in various fields. The enzyme’s efficiency and stability can be influenced by its structural components and the environmental conditions it encounters. Therefore, detailed structural studies of MDH can provide insights into its catalytic mechanisms and potential enhancements.^1-5^

We recently reported the inaugural crystal structure of pyrroloquinoline quinone (PQQ)-dependent methanol dehydrogenase (MDH) from the marine methylotrophic bacterium, *Methylophaga aminisulfidivorans* MP^T^ (MDH_Mas_), at a resolution of 1.7 Å.^1^ The active form of MDH_Mas_ (or MDHI_Mas_) is a heterotetrameric α_2_β_2_ complex, with each β-subunit symmetrically assembling on one side of each α-subunit (See Figure 1). This arrangement results in two β-subunits flanking the two PQQ-binding pockets on the α-subunits. Notably, the PQQ molecules are coordinated by a Mg^2+^ ion rather than the Ca^2□^ ion typically found in terrestrial MDHI, highlighting the enzyme’s ability to regulate osmotic balance in high-salt environments such as the ocean.

**Fig. 1.**
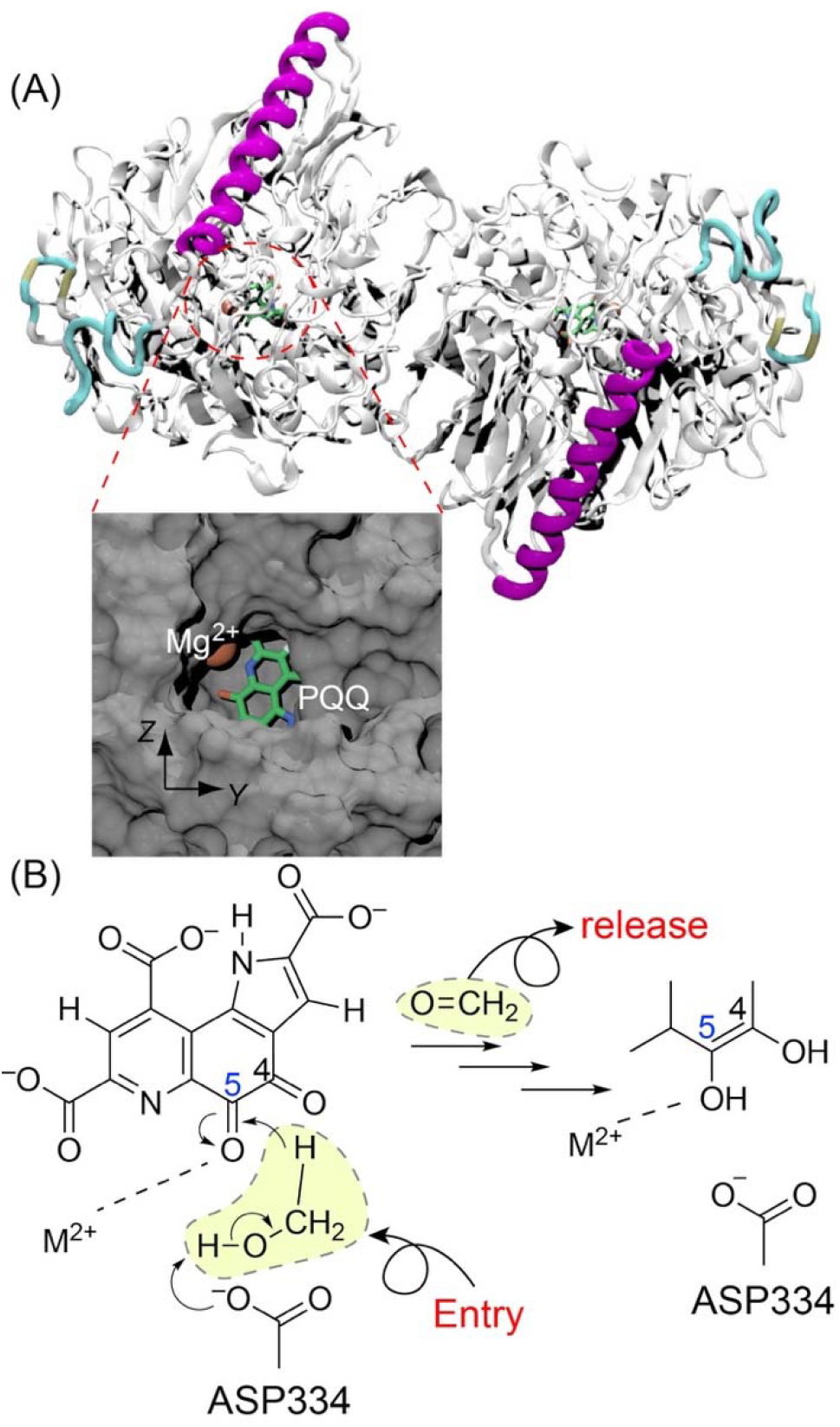
(A) Crystal structure of MDH from our previous report [PDB ID: 5XM3]. To represent the α_2_β_2_ structure, chains A and C are shown in a gray cartoon and chains B and D are shown in a colored cartoon model. Schematic representation of the active site containing PQQ and a Mg^2+^ ion is shown in inset. The plane of PQQ is defined as the yz-plane for our model. (B) Schematic representation of the enzyme activity mechanism of MDH_Mas_. MDH_Mas_ releases formaldehyde as a product after the oxidation of methanol. The full list of atom names, types, and partial charges of PQQ is provided in the supporting information (See Figures S1, S2, and Table 1).

However, even though we had a crystal structure of MDH_Mas_, it did not include the ligands such as methanol or formaldehyde, which is crucial for understanding the methanol dehydrogenase process. Therefore, we added the ligands to MDH_Mas_ by computational modeling and calculated the binding free energy of the ligands. We also performed MD simulations of the MDH_Mas_–ligand complex and analyzed the interactions between the ligand and MDH_Mas_. To the best of our knowledge, this is the first simulation that has determined the binding free energy between MDH_Mas_ and ligands at the atomistic level. This information is useful for the further design of more efficient MDH_Mas_ variants using protein design methods.^6,7^

Molecular dynamics simulations of proteins offer a comprehensive tool for studying the dynamic behavior of biomolecules. The ability to visualize and analyze protein motions at the atomic level provides profound insights into their function and interactions.^8-11^ We adapted the CHARMM36m force field for all of simulations.^12^ The detailed system is shown in Figure S4. One molecule of MDH was solvated in a water box of size 120 Å × 120 Å × 120 Å. This box was filled with 48,203 modified TIP3P water molecules, and 10 sodium ions were added to neutralize the system. An additional 136 sodium and chloride ions were added to maintain an ion concentration of 0.15 M.

To equilibrate the system, MD simulations were performed at 310 K for 1 ns with an NpT (constant number of particles, pressure, and temperature) ensemble following a 50,000-step geometry optimization. During equilibration, the positions of the oxygen atoms of crystal water molecules and the backbone atoms of MDH were constrained. A second equilibrium MD simulation was performed for 1 ns, constraining the positions of the Cα atoms of the backbone. The configuration obtained from this equilibration was used as the starting structure for the production simulation, which was conducted for 50 ns at 310 K with an NpT ensemble. The Langevin piston method maintained the pressure at 1 atm with a piston period of 100 fs, a damping time constant of 50 fs, and a piston temperature of 310 K.^13,14^ The full electrostatics were calculated using the particle-mesh Ewald method with a 1 Å grid width.^15^ The nonbonded interactions were calculated using a group-based cutoff with a switching function, updated every tenth time-step. The SHAKE algorithm was used to hold the covalent bonds involving hydrogen rigid,^16^ and a 2 fs time step was used. Atomic coordinates were saved every 100 ps for the trajectory analysis. The additional force field parameters for PQQ, methanol, and formaldehyde are shown in Figures S2 and S3, and Tables S1 and S2 in the supporting information. All MD simulations were carried out using nanoscale molecular dynamics (NAMD)^17^ and the graphics shown in this study were prepared using visual molecular dynamics (VMD).^18^

To evaluate the stability of MDH_Mas_ at 310 K, we performed a root mean square deviation (RMSD) analysis of the structures obtained from a 50 ns MD simulation. This measures the protein’s positional fluctuation relative to the crystal structure.

A snapshot of MDH_Mas_ after a 50 ns MD simulation is shown in Figure S5 (A), and the RMSD of chains A and B is shown in Figure S5 (B). The fluctuation of chains C and D are not shown because they are comparable to chains A and B. According to our analysis, the fluctuation of chain A is about 1 Å, and the fluctuation of the secondary structure is not significant (Figure S6 (A) in the supporting information), but the RMSD of chain B is slightly higher than that of chain A.

We calculated the fluctuation of each Cα atom of chain B and found that most of the fluctuation originates from the N-terminus of chain B, which exhibits turn and random coil secondary structures. The root mean square fluctuation (RMSF) of the Cα atom of chain B and a snapshot of chains A and B are shown in Figure S5 (C). The secondary structure of residues 25-35 in chain B fluctuates between the turn and random coil states (see Figure S6 (B) in the supporting information).

According to our previous crystallographic study, the active site of MDH_Mas_ is located on each α-subunit where the PQQ molecule is coordinated.^1^ The prosthetic group PQQ is structurally incorporated in an amphipathic way in the enzyme. PQQ resides within a hydrophobic pocket formed by the residues Cys134, Cys135, Val138, Trp274, Trp507, and Trp571. Additionally, PQQ is stabilized by hydrogen bonds with Glu86, Arg140, Thr190, Ser205, Glu208, Thr272, Asn425, and Trp507. After a 50 ns MD simulation at 310 K, we found that the position of PQQ is well maintained inside the hydrophobic pocket. Snapshots of PQQ obtained after the simulation are shown in Figure 2. During the simulation, the position of PQQ remained stable, and no significant translation was observed. However, C5 was exposed out of the hydrophobic pocket, despite PQQ being well-preserved inside the pocket through hydrogen bonding networks with neighboring residues.

**Fig. 2.**
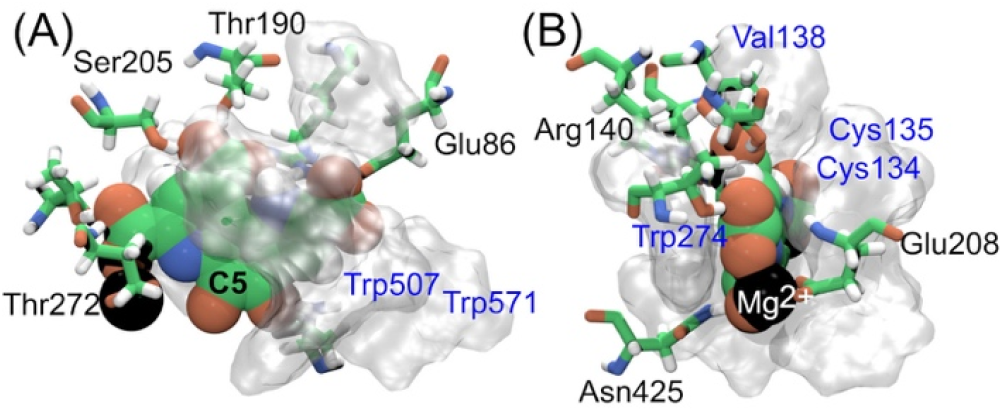
(A) Top and (B) side views of PQQ with neighboring residues. The residues forming the hydrophobic pocket for PQQ are represented by a transparent surface model, with each residue name shown in blue letters. The residues involved in the hydrogen bonding network with PQQ are shown in a licorice model, with each residue name in black letters.

The process of methanol dehydrogenase includes a general base facilitating proton removal from methanol, along with hydride ion transfer from the presumed methoxide to the quinone carbonyl carbon C5 of PQQ, followed by tautomerization to form hydroquinone PQQH_2_ (See Figure S1). Even though the ligand (methanol or formaldehyde) is absent in the crystallographic structure, we focus on methanol dehydrogenation process by MDH_Mas_. Therefore, we added ligands to the active site of MDH_Mas_ and studied the binding properties. Using the model complex, we calculated the activation free energy barrier of ligand binding (methanol or formaldehyde) in the active site of MDH_Mas_. Since we learned that the position of PQQ is stable during the MD simulation and the exit of the active site is normal to the PQQ plane, we defined the coordinates based on the position of PQQ. The plane of PQQ is taken as the yz plane, and the axis normal to the yz plane is taken as the x-axis. The ligand (methanol or formaldehyde) is pulled along the x-axis to calculate binding free energy.

We calculated the binding free energy of the ligand (methanol or formaldehyde) to MDH_Mas_, i.e., the potential of mean force (PMF), by pulling a ligand with a constant velocity (v_x_ = 1 Å/ns) using steered molecular dynamics (SMD)^19^ simulation from x_ligand_ = 0 to 10 Å (See Figure 3 (A)). The initial structure of MDH was adapted from the crystallographic structure in our previous report (PDB ID: 5XM3), and the ligand was placed at the binding site near PQQ. The system size and the conditions are the same as in the MD simulation.

**Fig. 3.**
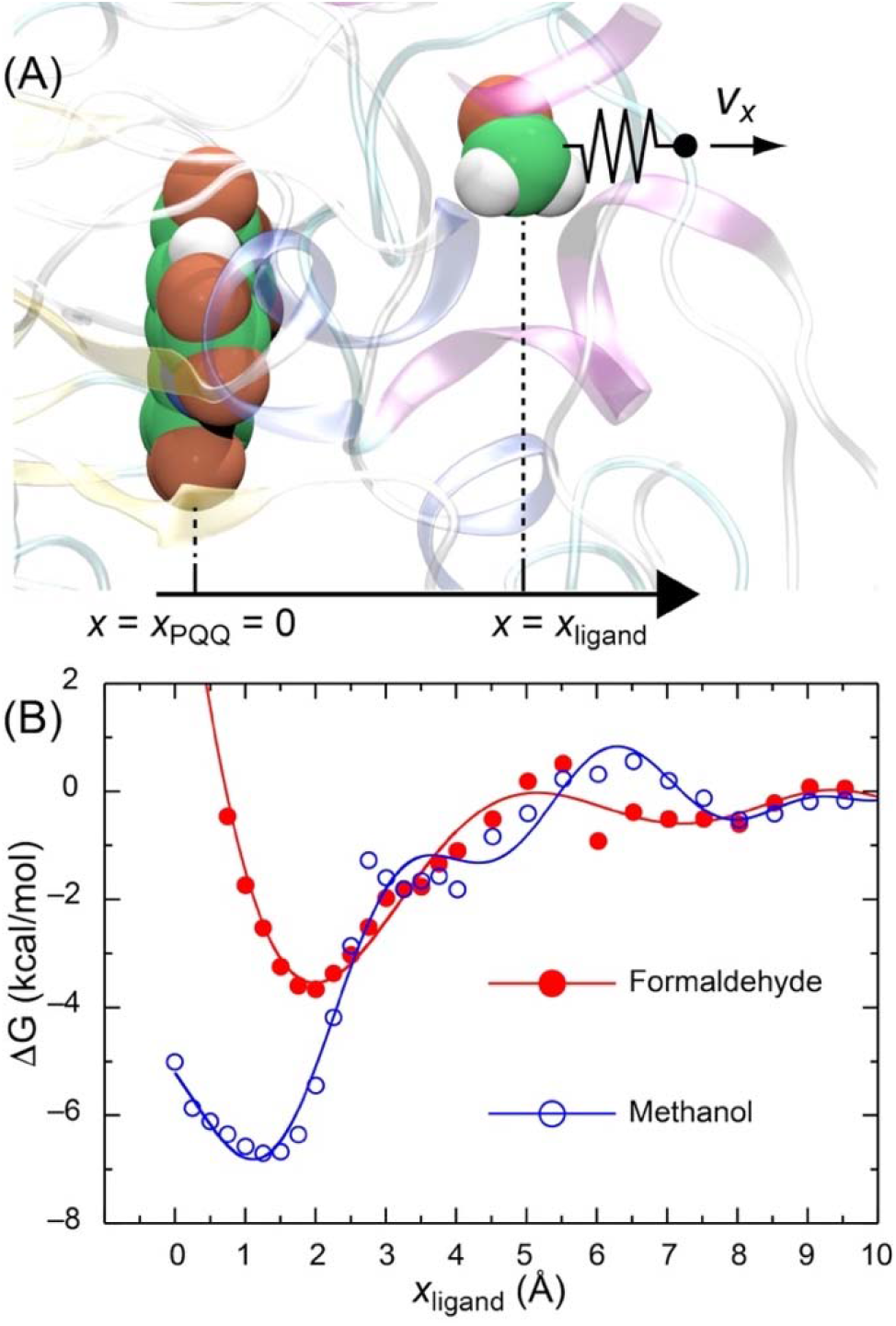
(A) Schematic representation of steered molecular dynamics (SMD) simulation. The plane of PQQ is adapted as the yz-plane, and the axis normal to the yz-plane is defined as the x-axis. The crystallographic position of PQQ is used for x_ligand_ = 0. The ligand is pulled at a constant velocity along the x-axis for the SMD simulation calculation. (B) Free energy profile of ligand binding to MDH_Mas_ obtained from SMD simulation. The binding free energy of methanol to MDH_Mas_ is about 7 kcal/mol, whereas the binding free energy of formaldehyde is about 4 kcal/mol.

The plane of PQQ is defined as the yz plane, and the axis parallel to the yz plane is used as the x-axis. The x-position of PQQ is set as x_ligand_ = 0. The path was divided into 10 smaller windows (each 1 Å in length), and 8 independent SMD simulations were performed in each window. The system was equilibrated in each window for 0.1 ns while constraining the x-position (defined by the positions of carbon and oxygen atoms only) of the ligand and the α-atoms of chain C of MDH at 310 K. The construction of the PMF from SMD simulations is based on the Jarzynski equality equation.^20^

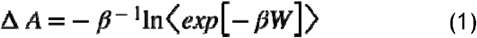

where ΔA is a free energy difference, ß is the product of the Boltzmann factor and temperature, and W is the non-equilibrium work obtained from SMD simulation. The non-equilibrium work done by the pulling force can be obtained using the following:

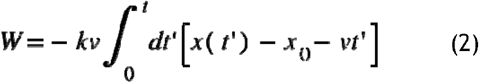

where k and v are the force constant (150 kcal/mol/Å^2^) and velocity of pulling (1 Å/ns), x(t’ s) and x_0_ are the reaction coordinate at t’ in the simulation, and the initial position of the center of mass of ligand. We adapted the second-order cumulant expansion equation for calculating Equation (2).

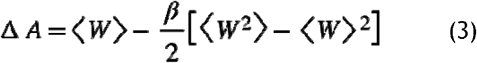

As expected from previous experimental suggestions, methanol shows the minimum free energy conformation around the C5 atom of PQQ when x_ligand_ = 1 Å (See Figures 3(B), 4, and Movie 1) for dehydrogenase reaction. When a methanol molecule approaches the active site of MDH_Mas_, it is near Arg130, Val137, and Phe578 when x_ligand_ = 10 Å.

As it approaches PQQ, methanol is near Cys135, Glu208, and Trp571 when x_ligand_ = 4 Å. The binding free energy of methanol is about 7 kcal/mol, whereas it is about 4 kcal/mol for formaldehyde. Therefore, we can expect that formaldehyde would be easily released from the active site after the methanol oxidation reaction. It appears that the entry path of methanol into the active site of MDH and the exit path of formaldehyde from the active site after the dehydrogenase reaction are similar. For example, both methanol and formaldehyde are in close contact with Trp571 at x_ligand_ = 5 Å, and their local minimum positions are very close: x_ligand_ = 2 Å for methanol and x_ligand_ = 1 Å for formaldehyde (see Figures 3 and 4).

**Fig. 4.**
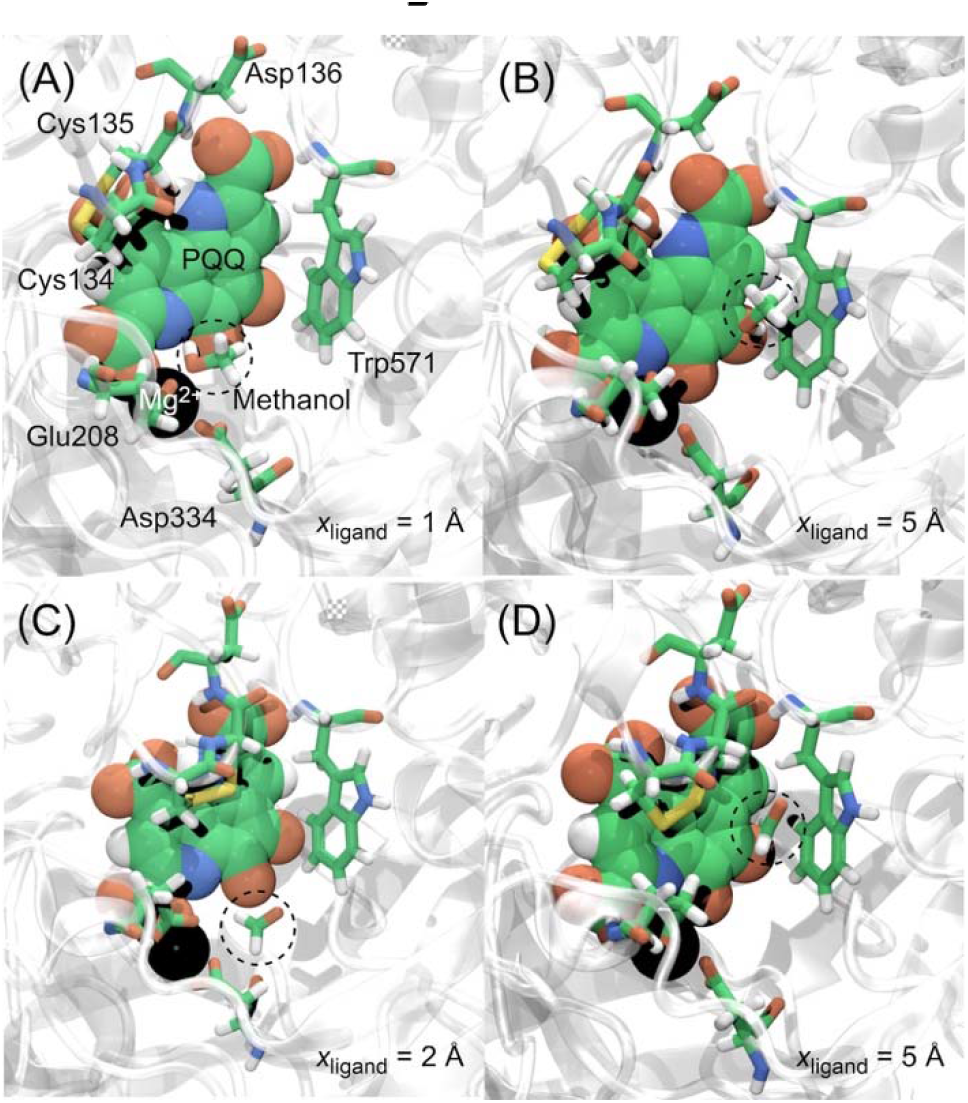
Snapshots of the binding complex structure of MDH with methanol at (A) x_ligand_ = 1 Å and (B) x_ligand_ = 5 Å; and with formaldehyde (C) x_ligand_ = 2 Å and (D) x_ligand_ = 5 Å.

The importance of protein design of MDH_Mas_ cannot be overstated. Through protein design, we can develop new MDH_Mas_ variants with higher efficiency of methanol dehydrogenase. Our current study provides valuable insights and could serve as a foundation for the further design and optimization of MDH_Mas_. By enhancing the catalytic properties and stability of MDH_Mas_, we can significantly improve its application in various industrial and biotechnological processes. As well as the simulated biophysical properties in the active site during catalysis, the information would help to simulate the docking location of the binding proteins such as Cyt c_L_^5^ and MxaJ,^2^ which are likely essential components oxidation.

## Supplementary data

Supplementary material Letters for complete methanol is available at Chemistry *Letters*

