## supporting information for "Computational Insights into the Molecular Dynamics of the Binding of Ligands in the Methanol Dehydrogenase"

Table S1. Atom name and the CHARMM atom type of PQQ. See Figure S2 for the name assignment.

| Atom name | CHARMM atom type | Atom name | CHARMM atom type |
| --- | --- | --- | --- |
| 1 | NG2R52 | 8 | CG2R63 |
| 1’ | HGP2 | 8’ | HGR62 |
| 1a | CG2RC0 | 9 | CG2R63 |
| 2 | CG2R51 | 9’ | CG2O3 |
| 2’ | CG2O3 | 9a | CG2RC0 |
| 3 | CG2R51 | O1 | OG2D2 |
| 3’ | HGR51 | O2 | OG2D2 |
| 3a | CG2RC0 | O3 | OG2D2 |
| 4 | CG2R63 | O4 | OG2D2 |
| 5 | CG2R63 | O5 | OG2D2 |
| 6 | NG2R60 | O6 | OG2D2 |
| 6a | CG2RC0 | O7 | OG2D2 |
| 7 | CG2R61 | O8 | OG2D2 |
| 7’ | CG2O3 |  |  |

Table S2. CHARMM atom type and partial charge of methanol and formaldehyde. The partial atomic charges were calculated with the B3LYP/6-31G* basis set using automated topology builder (ATB) server.^1^ See Figure S3 for atom number assignment.

| Methanol | | | Formaldehyde | | |
| --- | --- | --- | --- | --- | --- |
| Atom Number | Atom Type | Partial Charge | Atom Number | Atom Type | Partial Charge |
| 1 | CG331 | –0.04 | 1 | CG2O4 | 0.416 |
| 2 | OG331 | –0.65 | 2 | OG2D1 | –0.448 |
| 3 | HGA3 | 0.09 | 3 | HGR52 | 0.16 |
| 4 | HGA3 | 0.09 | 4 | HGR52 | 0.16 |
| 5 | HGA3 | 0.09 |  |  |  |
| 6 | HGP1 | 0.42 |  |  |  |

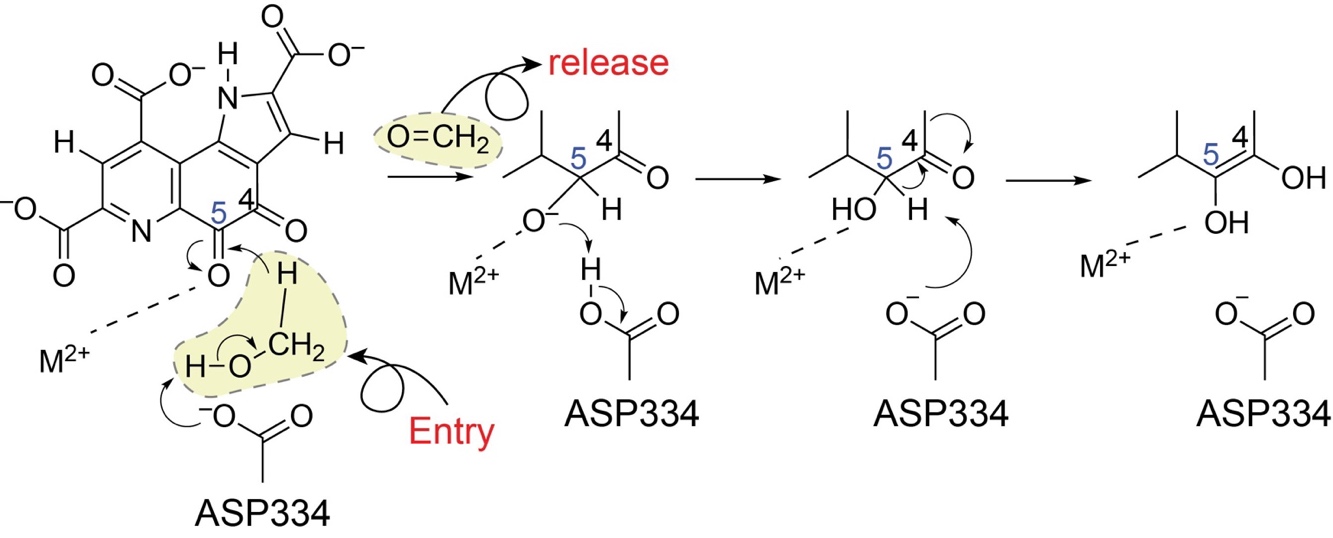

Figure S1. Schematic representation of the enzyme activity mechanism of MDH. MDH releases formaldehyde as a product after methanol dehydrogenation.

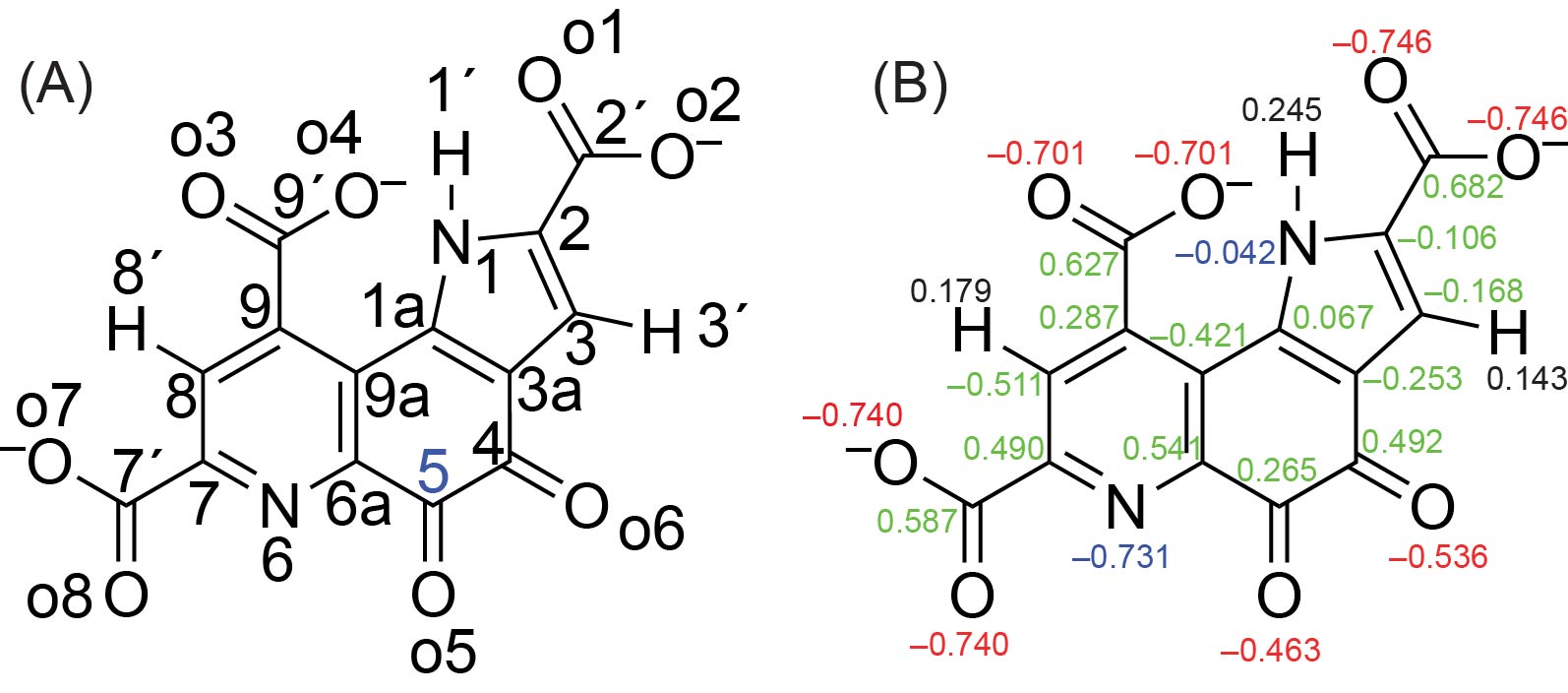

Figure S2. (A) Atom name and (B) atomic partial charge of PQQ. The partial atomic charges were calculated with the B3LYP/6-31G* basis set using automated topology builder (ATB) server.^1^

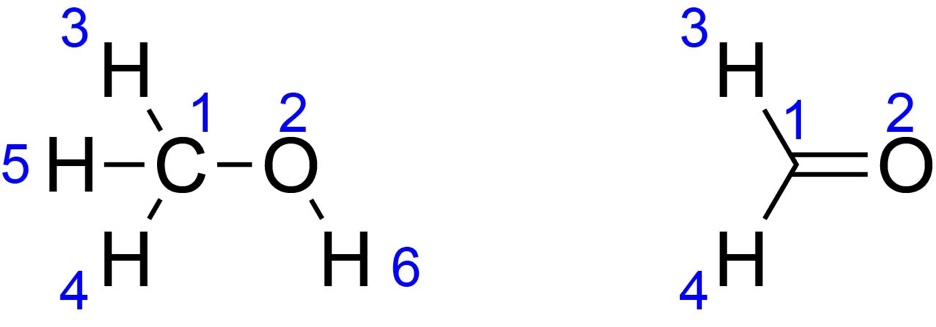

Figure S3. Atom number of methanol and formaldehyde used for current study

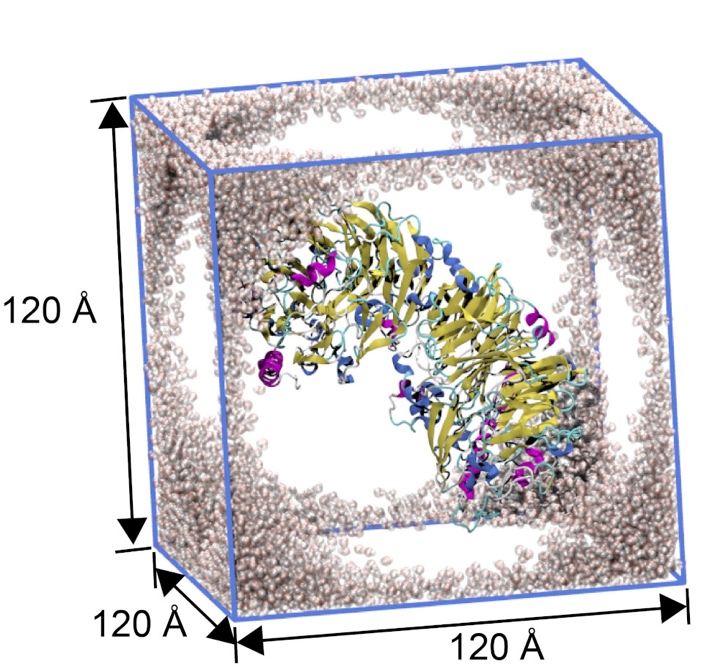

Figure S4. Schematic representation of the system. Only water molecules at the corners and edges are shown for clarity.

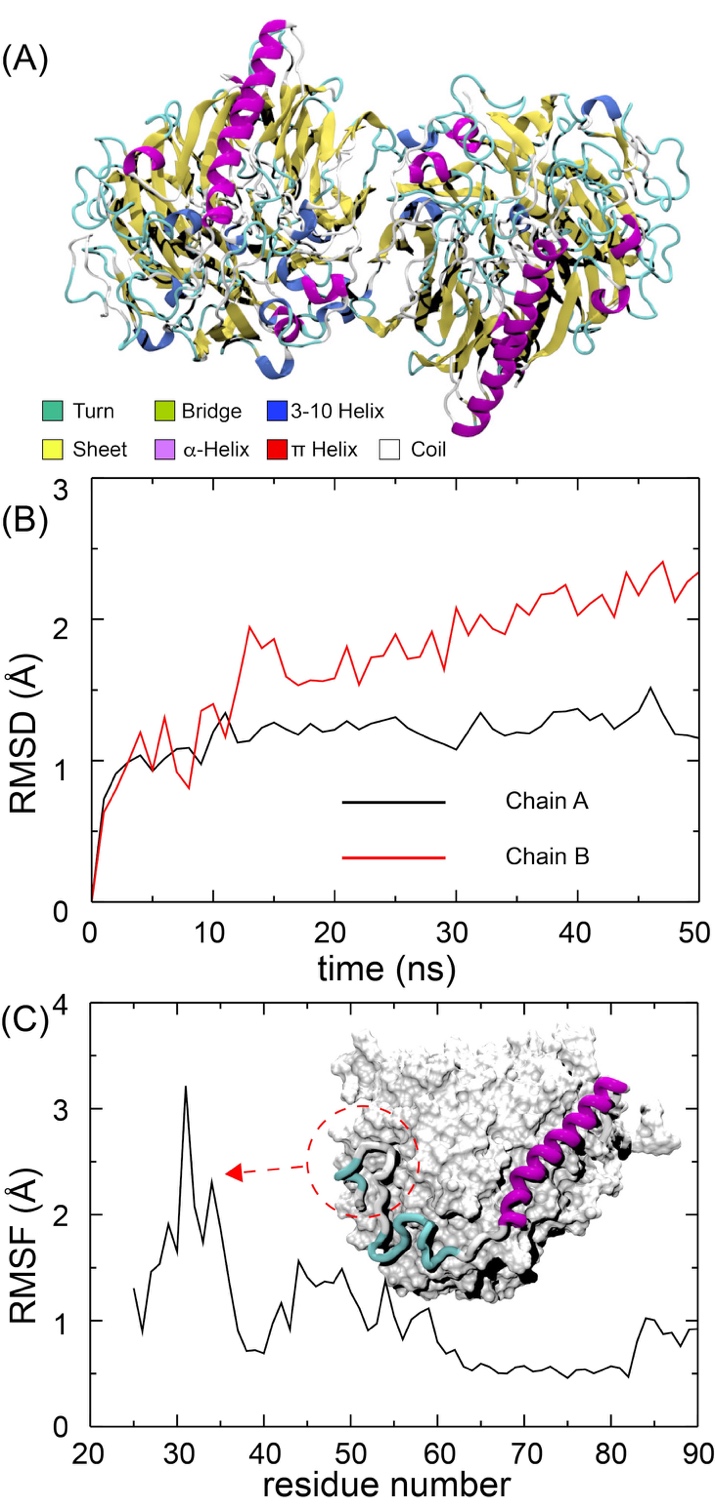

Figure S5. (A) Snapshot of MDH after 50 ns MD simulation, with color codes for the secondary structure of proteins. (B) Root mean square deviation (RMSD) of chains A and B during the MD simulation. (C) Root mean square fluctuation (RMSF) of Cα atoms in chain B over the 50 ns MD simulation. An inset shows a snapshot of chains A and B, with chain A represented by a surface model and chain B by a cartoon model.

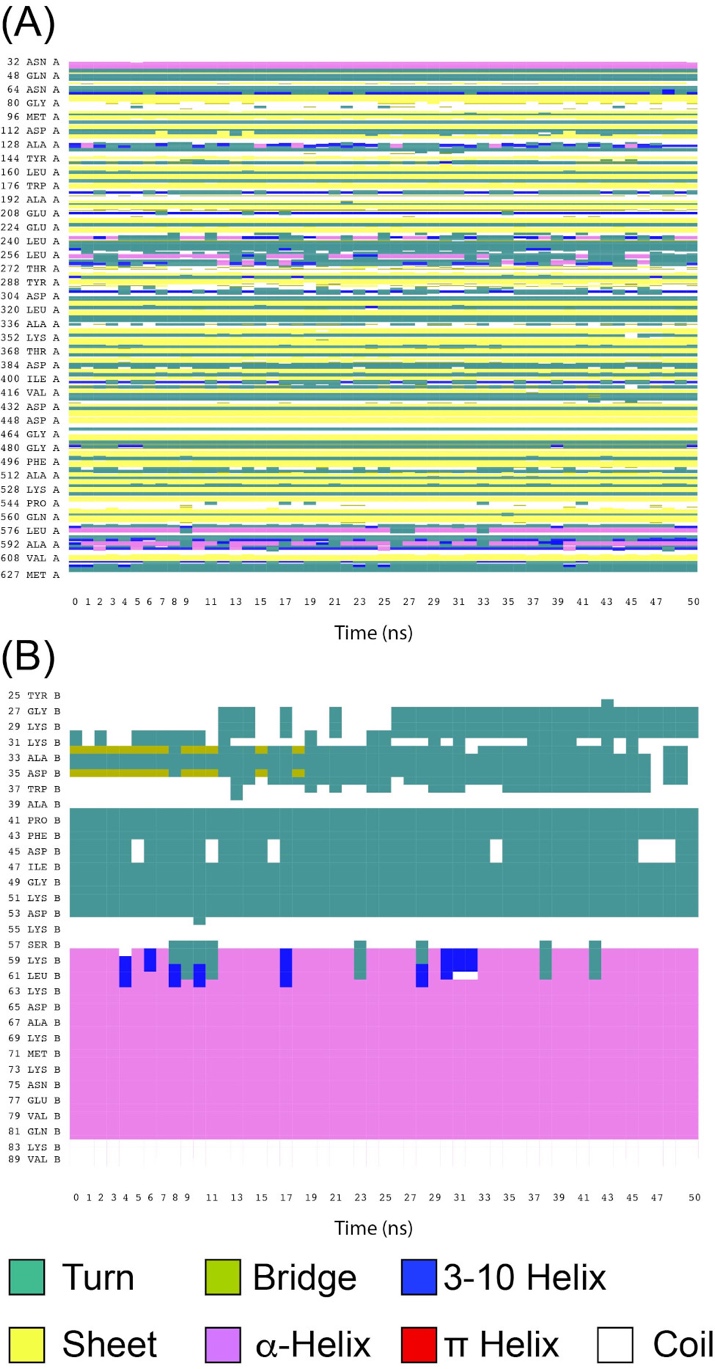

Figure S6. Secondary structure fluctuation of (A) chain A and (B) chain B during a 50 ns MD simulation, with color codes for each structure.
